# CryoEM Structures of Antibodies Elicited by Germline-Targeting HIV MPER Epitope-Scaffolds

**DOI:** 10.1101/2025.08.19.671101

**Authors:** Jiachen Huang, Olivia M. Swanson, Kimmo Rantalainen, Monica L. Fernández-Quintero, Johannes R. Loeffler, Ryan Tingle, Erik Georgeson, Nicole Phelps, Gabriel Ozorowski, Torben Schiffner, William R. Schief, Andrew B. Ward

**Author notes:** Correspondence (WRS), (ABW).

## Abstract

Applying cryoEM to small protein complexes is usually challenging due to their lack of features for particle alignment. Here, we characterized antibody responses to 21 kDa HIV membrane-proximal external region germline-targeting (MPER-GT) immunogens through cryoEM by complexing them with 10E8 or Fabs derived from MPER-GT immunized animals. Distinct antibody-antigen interactions were analyzed using atomic models generated from cryoEM maps. Mutagenesis screening revealed off-target mAbs that do not compete with 10E8 bind non-MPER epitopes, and the two most dominant epitopes were verified by cryoEM. The structures of 10E8-class on-target Fabs showed binding patterns that resemble the YxFW motif in the 10E8 HCDR3 loop. Additionally, we demonstrate that high-resolution maps can be generated from heterogeneous samples with pooled competing Fabs. Overall, our findings will facilitate the optimization of MPER GT-antigens and push the size limit for cryoEM-based epitope mapping with smaller antigens and heterogeneous antibody mixes.

**Highlights:**

- Elucidate the structural basis of increased 10E8 affinity to GT12 over GT10.2
- CryoEM epitope mapping reveals three main binding sites on MPER-GT immunogens
- Proof of concept for application of CryoEM epitope mapping to small immunogens
- Structural characterization and MD analysis confirms 10E8-like interactions of the MPER-specific Fabs

## INTRODUCTION

Inducing broadly neutralizing antibodies (bnAbs) remains a key objective in human immunodeficiency virus (HIV) vaccine development ^1^. A promising strategy to induce bnAbs through vaccination is germline-targeting (GT), which uses immunogens engineered to specifically activate rare naïve B cell precursors ^2–6^. The IAVI G001 phase I clinical trial (NCT03547245) demonstrated robust priming of B cell precursors of HIV VRC01-class bnAbs targeting the CD4 binding site using a germline-targeting immunogen, eOD-GT8 60mer ^7–10^. Similar immunogens are being developed to target other Env broadly neutralizing epitopes, including the V3-glycan^11–14^, V2-apex ^15,16^, and the fusion peptide ^17^. Several of these immunogens have been shown to prime bNAb precursors in animal models ^18–20^.

The highly conserved Env membrane proximal external region (MPER) epitope is another attractive target for germline-targeting vaccine design, as MPER-specific bnAbs, including 10E8 ^21^, LN01 ^22^, DH511 ^23^, and 4E10 ^24^ exhibit high neutralization coverage across diverse HIV strains. However, the MPER epitope is usually absent from engineered soluble recombinant Env immunogens, since it reduces the stability and solubility of the Env trimer ^25^. Its recessed location at the base of the Env trimer also limits the access for naïve B cells. To present the MPER epitope in soluble form for vaccine delivery, the MPER helix was grafted onto engineered protein scaffolds, with additional modifications to improve the affinity for 10E8-class precursors, resulting in a series of germline targeting immunogens termed 10E8-GT ^26^. Multivalent nanoparticles composed of 10E8-GT scaffolds displaying germline-targeting MPER epitopes, such as 10E8-GT10.2 12mer and 10E8-GT12 24mer, successfully primed rare 10E8-class bnAb-precursor responses in transgenic mouse models and non-human primates (NHPs) ^27,28^. While both 10E8-GT10.2 and 10E8-GT12 primed comparable levels of 10E8-class responses in vivo, 10E8-GT12 had much higher affinity for mature 10E8-class variants and induced more key 10E8-class mutations in primed 10E8-class B cells ^27^. Structures of 10E8-GT immunogen in complex with 10E8-class antibodies showed conserved interactions of the targeted HCDR3 “YxFW” motif, but a range of different binding angles were discovered, posing additional questions regarding the differences of binding modes ^27^. Although most of these structures were determined by X-ray crystallography, some complexes were difficult to crystallize even after extensive condition screening. Moreover, inaccuracies in model building could be introduced due to crystal packing, which might lead to misinterpretation of the structural models.

Electron microscopy (EM)-based polyclonal epitope mapping (EMPEM) can be used to characterize humoral immune responses after infections or vaccinations, offering structural insights on antibody-antigen interactions at both low and high resolution levels ^29^. While EM-based epitope mapping has been successfully applied to trimeric viral fusion proteins such as influenza hemagglutinin (HA), SARS-CoV2 spike, and HIV Env ^30–34^, its application to small proteins remains challenging, and often requires complexation with other proteins increase the overall complex mass, thereby facilitating particle alignment ^35^. A recently published CryoEM structure of 10E8-GT10.2 ^27^, demonstrated that it is feasible to apply cryoEM-based epitope mapping on small asymmetric protein complexes like MPER-GT immunogens when complexed with two or more Fabs.

Here, we further characterized the structures of 10E8-GT12 and the monoclonal antibodies induced by the 10E8-GT10.2 and 10E8-GT12 immunogens (referred as GT10.2 and GT12 below) using cryoEM. In addition, we explored the application of cryoEM to ∼120 kDa, heterogeneous protein complexes (with ∼70 kDa models resolved with Fab variable regions) by mixing multiple competing Fabs in the same sample. Despite limitations due to preferred orientation and the inherent challenges of cryoEM data processing for small particles, we managed to identify specific Fab densities from a pool of three MPER-targeting Fabs, and generated ∼ 4 Å 3D reconstruction maps revealing distinct binding patterns of MPER-targeting antibodies. Molecular dynamics (MD) simulations further revealed the flexibility of the Fab CDR loops when interacting with the MPER epitope. In summary, our analysis provides structural insights into10E8-like, and off-target mAbs induced from animals immunized with GT10.2 12mer and GT12 12mer. These findings are crucial for understanding how these antibodies are generated and how to optimize boosting immunogens to more effectively induce neutralizing antibodies targeting full-length Env.

## RESULTS

### 10E8 binds GT12 in a similar manner compared to previously reported MPER complexes

Mature 10E8 has been shown to bind 10E8-GT10.2 (PDB 8SX3, Fig. 1A) in a manner that mimics previously reported crystal structures of the MPER peptide (PDB 4G6F) ^21^ and non-germline-targeting epitope scaffold T117v2 (PDB 5T85) ^26^, with almost identical interactions of the HCDR3 YxFW motif. The partially germline-reverted 10E8-class antibody 10E8-iGL1 also bound with similar binding modes to 10E8-GT4 (PDB 8TZW), and 10E8-GT11 (PDB 8U08) ^27^. However, structural evidence verifying 10E8 binding to GT12 is still lacking. Here, we determined a 4.1 Å cryoEM structure of 10E8-GT12 in complex with mature 10E8 Fab and the previously reported off-target mouse Fab W6-10 ^27^ (Fig. 1B, Fig. S1A&B). An overlay of the Fv regions and HCDR3 loops of 10E8 demonstrated its similar binding angles to GT10.2 and GT12, with a root-mean-square deviation (RMSD) of 0.77 Å. The HCDR3 sidechains, especially the D gene-encoded YxFW motif, overlapped uniformly between GT10.2+10E8 and MPER peptide+10E8 complexes (Fig. 1C). Based on GT10.2, the MPER helix residues around the binding pocket of GT12 were redesigned to facilitate higher 10E8 affinity (Fig. 1D) ^27^. While most of the key residues were retained in GT12, the mutations introduced in GT12 altered the surface hydrophobicity landscape of the binding pocket (Fig. 1E) and might have increased flexibility of the MPER helix, as suggested by molecular dynamics analysis (Fig. 1F, Video 1 & 2). Further analysis of interactions between the epitope and paratope revealed that Y79, W98, and K109 in GT10.2 potentially form hydrogen bonds (H-bonds) with multiple sidechain and backbone atoms of the 10E8 HCDR3, while residues L42, W98, V102, I103, and L105 mediate hydrophobic interactions with the 10E8 YxFW motif (Fig. 1G). In GT12, mutations V102T and I103N introduced H-bonds with the backbone of 10E8 HC P100g, G100h, and Y99, respectively. These additional interactions likely contribute to the increased affinity of GT12 to 10E8. Beyond the interactions between the MPER epitope and 10E8 HCDR3 loop, HCDR1 W33, HCDR2 E53, and HCDR3 K97 form a pocket that sequesters the turning point (W98) of MPER helix (Fig. S1C), similar to the previously reported structure of 10E8 Fab-MPER peptide complex (PDB 4G6F). Finally, in both GT10.2 and GT12 complexes, D99 forms a salt bridge with 10E8 LC R95b, further strengthening the interaction between 10E8 and the GT scaffold (Fig. S1C), whereas residue F95b in native gp41 is unlikely to mediate any interaction with the 10E8 LC.

**Figure 1.**
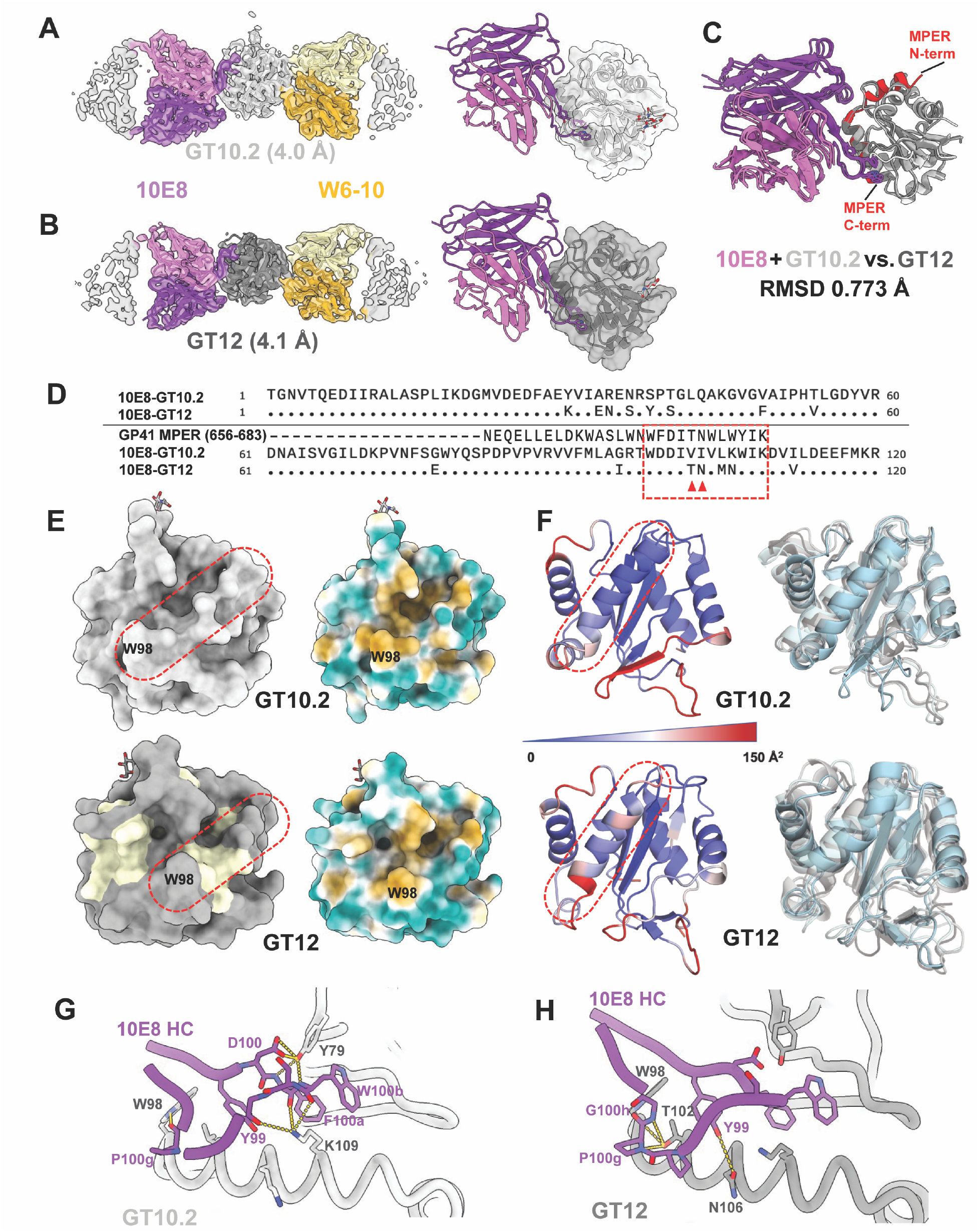
10E8 shares similar binding interactions with GT10.2 and GT12. CryoEM maps and models of 10E8 (purple)/W6-10 (yellow) Fabs complexing with GT10.2 (A) and GT12 (B). Aligned structures (C) and sequences (D) of 10E8 Fab with GT10.2, GT12, and the MPER peptide (red, PDB: 4G6F). (E). The designed MPER helix is circled in red dash boxes, the engineered GT12 residues that are different from GT10.2 are colored in light yellow (left panel). Surface hydrophobicity analysis of GT10.2 and GT12, the hydrophobic sidechains are colored in orange while hydrophilic sidechains were colored in teal (right panel). (F). GT10.2 and GT12 scaffolds color coded by the B-factor obtained from MD simulations as a measure of flexibility to identify regions with higher (red) and lower (blue) variability (left panel), with the most frequent states from MD analysis are overlayed (right panel). The residues involved in the interactions of 10E8 HCDR3 loop with GT10.2 (G) and GT12 (H) are displayed.

### A GT12-induced 10E8-like mouse antibody mimics 10E8 binding to GT12

10E8-class monoclonal antibodies were previously generated from antigen-specific B cells isolated after 10E8-GT12 12mer vaccination from mice engineered to produce antibodies with human D_H_3-3 and J_H_6 segments ^27^. Epitope footprints for recombinantly expressed IgGs/Fabs were mapped by binning assays. Next, we complexed the GT12-induced 10E8-class Fab, SA2911, with GT12 as well as the off-target Fab W6-10, and a 4.0 Å cryoEM structure was obtained (Fig. 2A, Fig. S1D & E). SA2911 shares similar HCDR3 interactions with 10E8 in the GT12 MPER binding pocket (Fig. 2B). Both heavy and light chains (HC and LC) of SA2911 are involved in interactions with GT12. HCDR3 Y98 and Y99 potentially form H-bonds with GT12 H53 and W98, respectively. HCDR3 Y100g occupies the binding pocket through interactions with GT12 N106 and K109 (Fig. 2C). The SA2911 light chain contains a long LCDR1 loop (17aa), where N28 and Q29 interacted with GT12 Y79. Together with LCDR1, LCDR2 Y49, F50, and R54 form a binding pocket that engages the GT12 77-81 loop (Fig 2D). These additional LC interactions likely contribute to the shift of binding angle of SA2911 by 20.8° relative to 10E8 (Fig. 2B).

**Figure 2.**
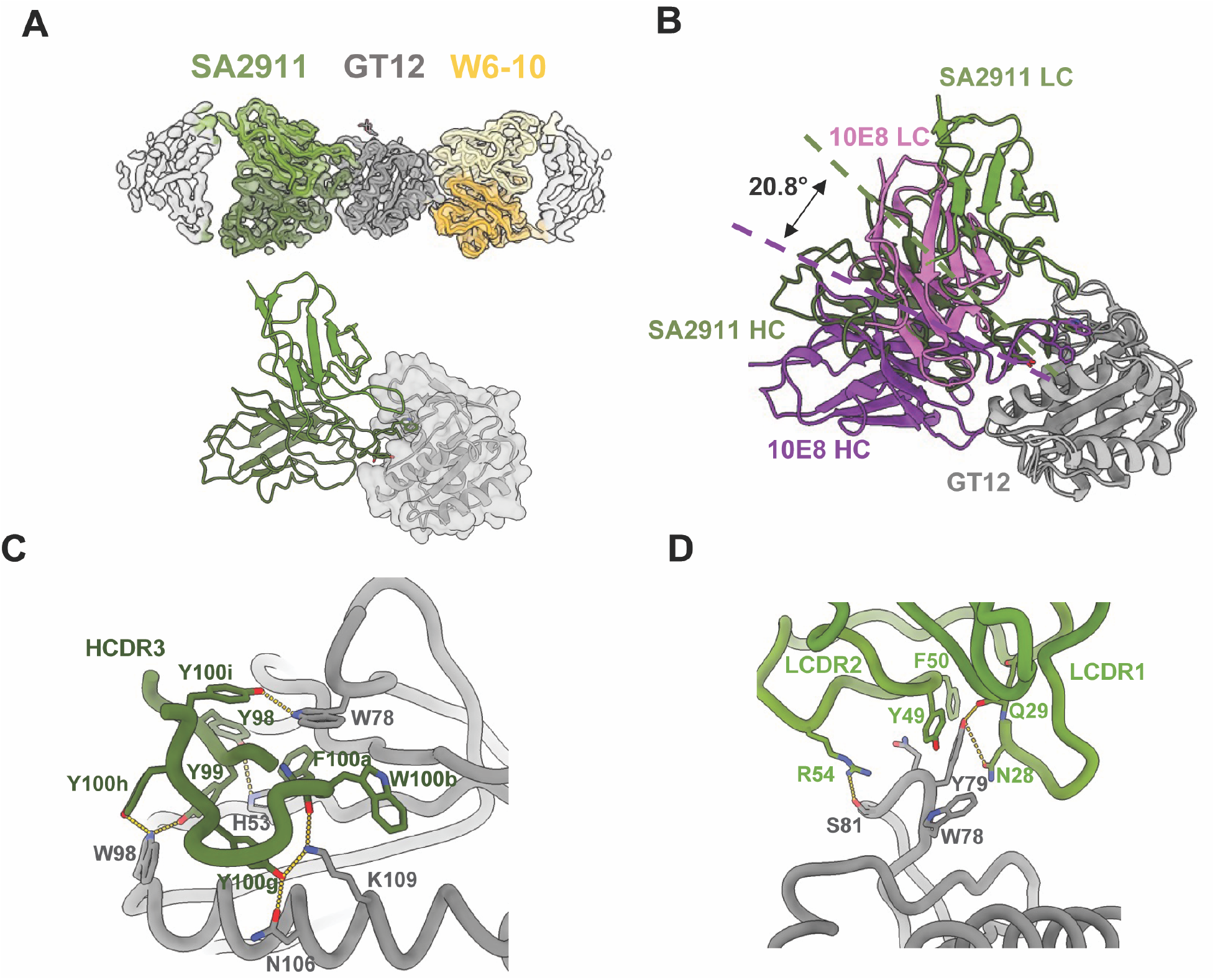
GT12-induced antibody SA2911 showed 10E8-like binding patterns on the MPER helix. (A) CryoEM map and model of SA2911 (green)/W6-10 (yellow) Fabs complexing with GT12. (B) Aligned structures of 10E8 (purple) and SA2911 (green) Fabs on GT12, with dashed lines showing different angles of approach. The angles are measured between mAb centroids and the GT12 centroid. The residues that interact with GT12 in W3-05 Fab HC (C) and LC (D) are displayed.

### Structural determination of Fab W3-05 from a pool of three 10E8-class mAbs complexed with GT10.2

We next set out to provide a proof-of-concept for applying cryoEM epitope mapping ^36,37^ to smaller antigens like the MPER-GT scaffolds. Three 10E8-class on-target Fabs (W3-02, W3-05 and W10-09) and one off-target Fab (W6-03) were pooled and complexed with GT10.2 at equimolar ratios of Fabs:off-target Fab:GT10.2 antigen for data collection (Fig. S1F). After initial particle selection through 2D classifications, particles were further classified by ab-initio reconstructions, and the volumes with clear Fab densities were selected. After three rounds of selection, only two classes were resolved with good structural details, while the third class failed to reach interpretable resolution (Fig. S2A). The second round of refinement of the two best classes resulted in 3.9 Å and 4.2 Å resolution reconstructions. The densities of on-target Fab between the two classes could be distinguished by their binding angles (Fig. S2B). Among the on-target Fabs, the W3-05 sequence best fit the 3.9 Å map (Fig. 3A, Fig S1G), while the 4.2 Å map lacked sufficient sidechain density to determine its identity due to its lower resolution. To resolve this ambiguity, we collected a separate dataset for GT10.2 complexing with W10-09 Fab and W6-10 Fab, resulting in a 6.3 Å reconstruction that clearly showed a different binding angle of W10-09 compared to the two maps determined earlier (Fig. S2C & D), suggesting that the 4.2 Å map likely corresponds to W3-02. W10-09 Fab also had the lowest binding affinity among the pooled Fabs (Table 1), indicating that W3-02 and W3-05 Fabs might have outcompeted W10-09 Fab when complexed together. Compared to 10E8, W3-05 has flipped HC/LC orientations and a 14° shift in binding angle, while the YxFW motif of the HCDR3 retained key interaction with the GT10.2 binding pocket (Fig. 3B). Potential H-bonds were observed between HC Y99 and GT10.2 R34/H58, as well as HC N100 and GT10.2 T40/G41 (Fig. 3C). In addition, residues K52, D56, and T59 in the HCDR2, K31 and Y32 in LCDR1, and D95a in LCDR3 interact with GT10.2 residues outside the binding pocket (Fig. 3D & E).

**Table 1.** Summary of Fab affinities to MPER scaffolds.

| Fab | Alias | Source | Target | GT10.2 kD (nM) | GT12 kD (nM) |
| --- | --- | --- | --- | --- | --- |
| 10E8 | - | - | MPER | 247 | 1 |
| NHP-10E8_MPER-GT10v55_W3-05 | W3-05 | GT10.2 immunized NHP | MPER | 19 | 490 |
| NHP-10E8_MPER-GT10v55_W10-09 | W10-09 | GT10.2 immunized NHP | MPER | 30 | 890 |
| NHP-10E8_MPER-GT10v55_W3-02 | W3-02 | GT10.2 immunized NHP | MPER | 7.5 | 3400 |
| NHP-10E8_MPER-GT10v55_W10-08 | W10-08 | GT10.2 immunized NHP | MPER | 45 | 29 |
| NHP-10E8_MPER-GT10v55_W10-07 | W10-07 | GT10.2 immunized NHP | MPER | 3.8 | 110 |
| NHP-10E8_MPER-GT10v55_W3-08 | W3-08 | GT10.2 immunized NHP | MPER | 6.2 | 74 |
| SA2102_10E8-like_2911 Fab | SA2911 | GT12 immunized mice | MPER | 787 | 534 |
| SA684_Exp1_Scaf_MPER-GT10v55_W6-03 | W6-03 | GT10.2 immunized mice | Off-target | 9 | 8.5 |
| SA684_Exp1_Scaf_MPER-GT10v55_W6-01 | W6-01 | GT10.2 immunized mice | Off-target | 9 | 0.84 |
| SA684_Exp1_Scaf_MPER-GT12v56_W6-10 | W6-10 | GT12 immunized mice | Off-target | 3 | 1.8 |

**Figure 3.**
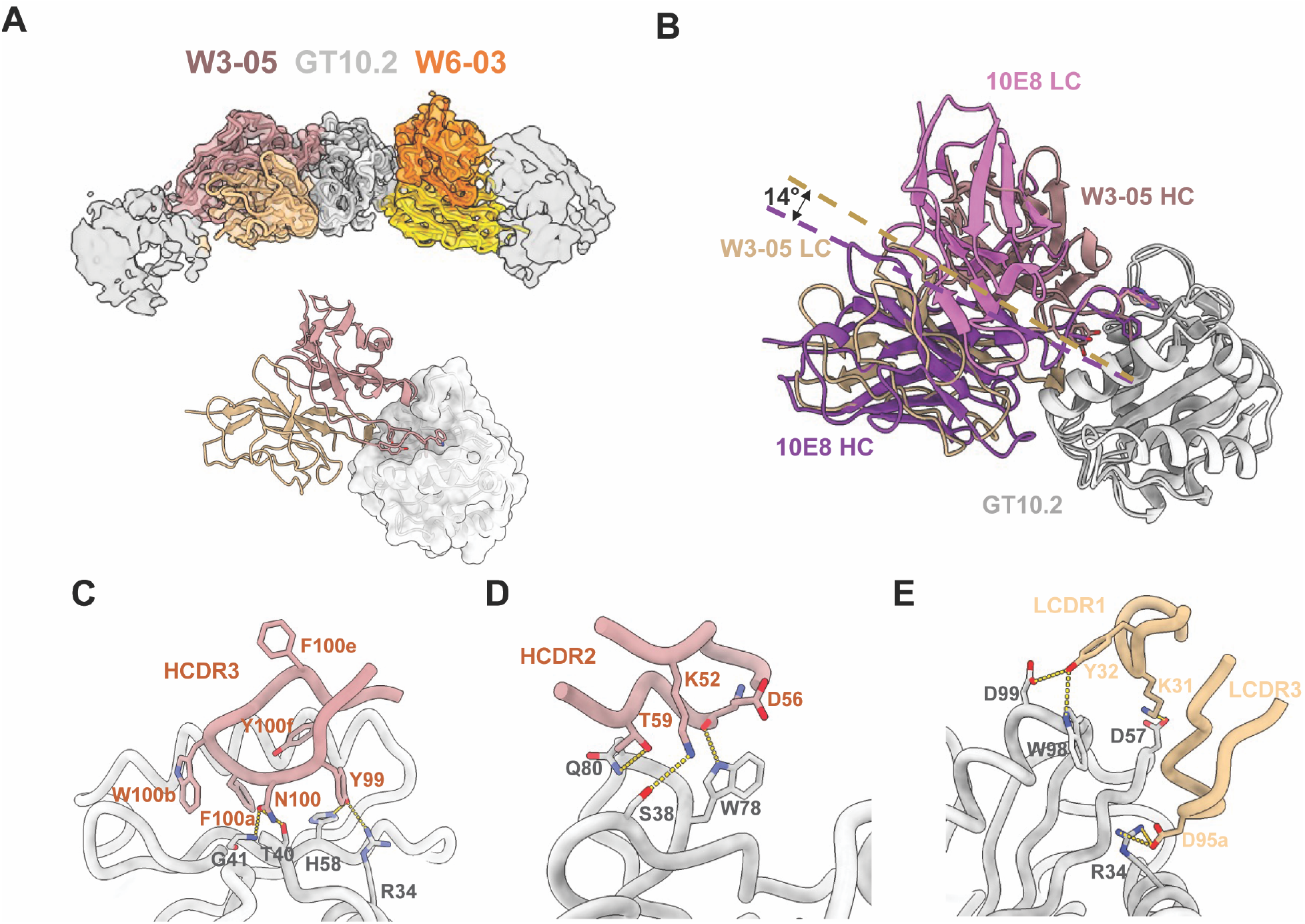
W3-05 binds to GT10.2 with flipped HC/LC positions comparing to 10E8. (A). CryoEM map and model of W3-05 (brown)/W6-03 (orange). (B) Binding angle shift revealed by aligning GT10.2 complexes with W3-05 Fab vs. 10E8 Fab, the angles are measured between mAb centroids and the GT12 centroid. The residues that interact with GT10.2 in W3-05 Fab HCDR3 (C), HCDR2 (D), and LC (E) are displayed.

To further increase sample heterogeneity to mimic the polyclonal composition in the serum, we complexed GT10.2 with a pool of six 10E8-class on-target Fabs and three off-target Fabs and collected cryoEM data. Aligned particles from 2D classification revealed a new off-target binding site oriented perpendicularly to the MPER epitope and the off-target epitope described above (Fig. S3A), consistent with antibody binning results that indicated the epitope for Fab W6-01 was different from W6-03 and W6-10 (Fig. 4A &B, Table S2). Densities of the on-target Fabs in the 2D classes suggested engagement of the MPER with a variety of angles of approach (Fig. S3B). Unfortunately, this dataset suffered from severe preferred orientation (Fig. S3C), and was unsuitable for further data processing and 3D reconstruction. We hypothesized that W6-01 being on the same plane with the other two binding sites limits tumbling of the complexes.

**Figure 4.**
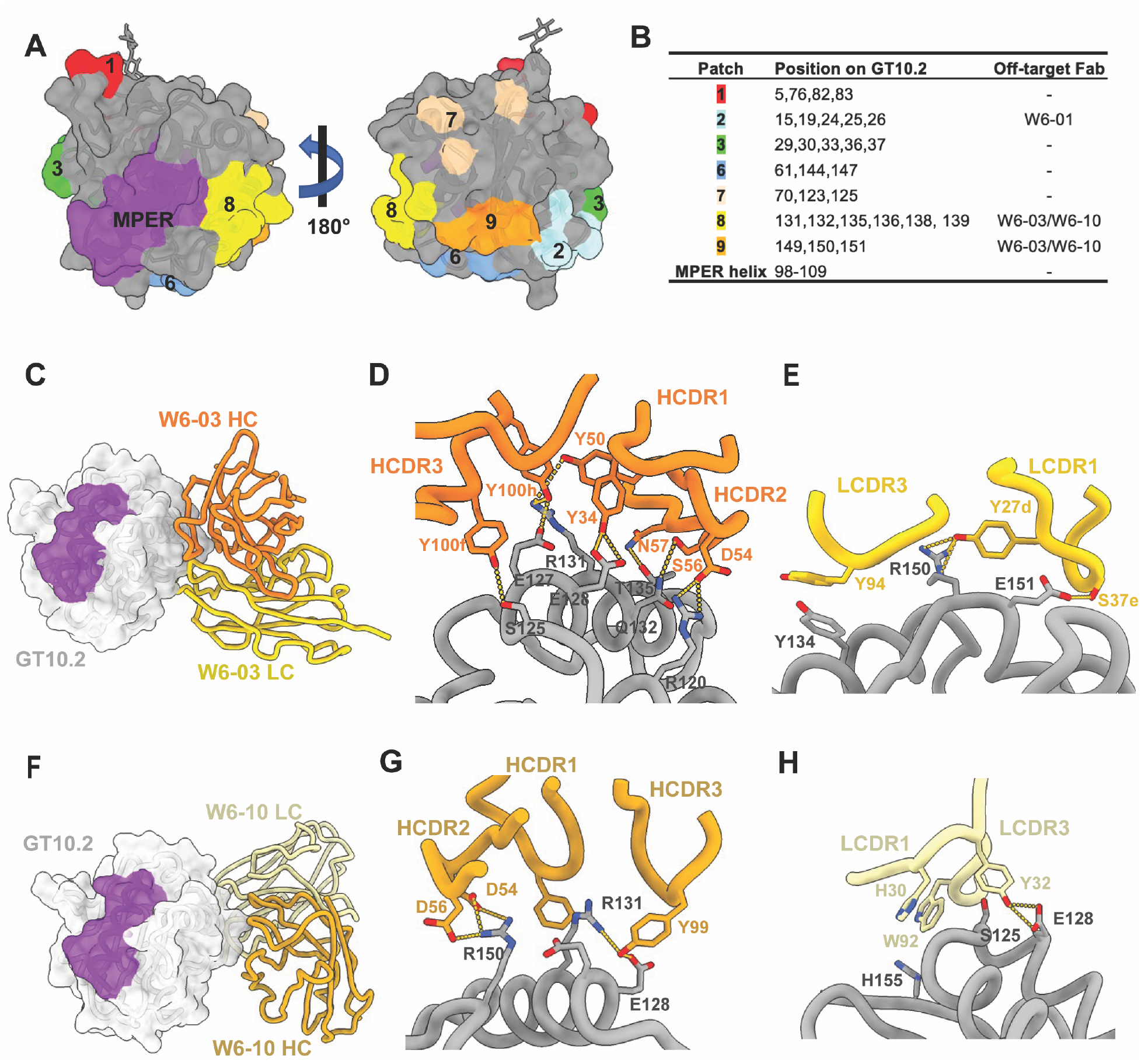
Epitope mapping of off-target Fabs and analysis of their interactions with GT-scaffolds. The off-target patch mutation residues are displayed in (A) and the epitope mapping of off-target fabs are listed in (B). The binding angles and heavy/light chain interactions of W6-03 (C-E) and W6-10 (F-H) are displayed.

### Structural comparisons of off-target Fabs

The cryoEM datasets described above enabled us to resolve structures of three antibodies recognizing two distinct epitopes on GT10.2 (Fig. 4A & B). The epitopes of the off-target Fabs W6-03 (induced by GT10.2 12 mer) and W6-10 (induced by GT12 12 mer) were highly similar in sequence between GT10 and GT12, and the binding affinities of the two mAb-scaffold complexes were comparable (Table 1). W6-03 and W6-10 have overlapping footprints but slightly different angles of approach (Fig. 4C & F). Both HC and LC of the off-target antibodies mainly interact with charged residues on the scaffold, including E128, R131, and R150 (Fig. 4D-E & G-H). In addition to the three off-target Fabs characterized above, the binding sites of 16 additional off-target antibodies were mapped with the GT10.2 patch mutants, and five of them showed similar binding profiles with patches 2, 8, and 9 (Table S2). This suggests that these off-target epitopes are immunogenic and modifications of these residues would be beneficial to mitigate off-target responses against these regions.

### Structural comparisons of 10E8-class Fabs

Overall, the on-target Fabs characterized above showed 10E8-like binding patterns, especially with the HDCR3 loops, yet with notable differences (Fig. 5A). While the 10E8 LC barely interacts with MPER-GT immunogens, both HC and LC of W3-05 and SA2911 formed interactions with the MPER binding pocket as well as the scaffolds. In GT10.2, R34 and H53 form H-bonds with HCDR3 Y99 (the first tyrosine in the YxFW motif) that pull the Y99 away from the hydrophobic binding pocket, and Y100e takes the position within the binding pocket (Fig. 5C). A similar “tyrosine switch” was observed with SA2911 bound to GT12, where Y100g occupies the pocket (Fig. 5D), as well as bnAb LN01 (F100f) ^22^ and the previously reported W3-01 (Y100f) ^27^ isolated from GT10.2 immunized macaques (Fig. 5E). To further understand how the “tyrosine switch” affects the antibody binding angle and affinity, we performed MD simulation analysis with W3-05 and SA2911 to inspect potential dynamics of HCDR3 sidechains, especially the aromatic residues, within the binding pocket. While W3-05 did not show significant sidechain movements compared to 10E8, SA2911 exhibited substantial instability as the conformation of HCDR3 backbone and sidechains were significantly rearranged during the simulation (Fig. 5F & G, Video 3 & 4). This might have been due to the lower affinity of SA2911 against MPER-GT antigens compared to W3-05 (Table 1), and suggested that the “tyrosine switch” can be acommodated by flexible HCDR3 conformations.

**Figure 5.**
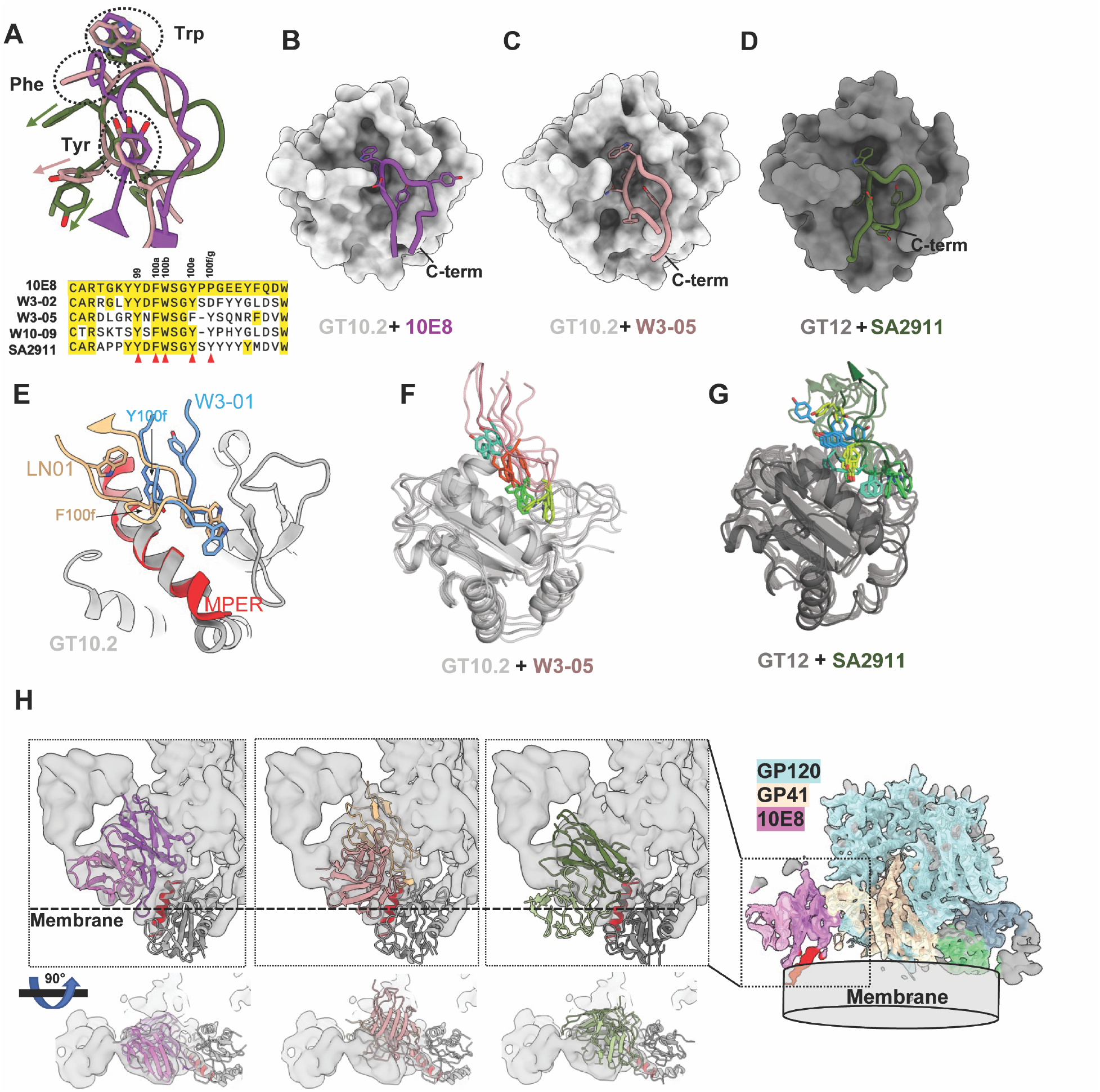
GT10.2/GT12 primed 10E8-class antibodies have distinct binding angles and HCDR3 conformations in the binding pockets. (A). The HCDR3 loops are aligned with clustered side chains of YxFW motif in the dashed circles. The layouts of HCDR3 loops of 10E8 (B), W3-05 (C), and SA2911 (D) are shown on the surface of GT10.2/GT12. (E). The HCDR3 loops of LN01 and W3-01 are aligned on GT10.2 binding pocket. Representative states of W3-05 (F) and SA2911 (G) HCDR3 loops generated by MD simulation. (H). The on-target fab-GT complexes are aligned with the MPER peptide and cryoEM map of membrane-bound full-length Env (PDB:6VPX, EMD-21335), the density of the nanodisc is indicated as membrane.

10E8 Fab is able to access the MPER epitope of full-length Env within the lipid bilayer ^25^, which is facilitated by interactions of 10E8 LC with lipid head groups ^26,38–40^. To understand how the binding angles of 10E8-like mAbs correlate with potential membrane interactions, we manually docked the structures of W3-05 Fab and SA2911 Fab into a previously reported cryoEM map of full-length Env. The distance between the membrane density and W3-05 HC is similar to that of 10E8 LC, but W3-05 LC clashes with gp41 density, suggesting W3-05 might be unable to bind full-length Env. On the other hand, SA2911 does not clash with any part of Env, however, its LC appears to be buried deeper within the membrane lipid density, which may prevent it from binding to full-length Env (Fig. 5H). We note that neither of these Fabs were elicited by vaccination regimens that included a membrane-bound Env immunogen, but that fuller regimens including membrane-bound Env have been proposed as necessary to drive complete maturation to bnAb ^27^. These observations imply that binding to full-length Env imposes strict geometric constrains, and further underscore the importance of precisely guiding antibody maturation through boosting immunogens to achieve compatible binding angles.

## DISCUSSION

In this study, we analyzed the structural basis of antibody responses to MPER-GT immunogens by determining cryoEM structures of GT10.2 and GT12 in complex with 10E8 and on/off-target antibodies derived from vaccinated animals. We verified that mature 10E8 binds to GT12 in a manner consistent with its binding to MPER peptide and previous versions of MPER-GT immunogens and elucidated the molecular basis of increased 10E8 affinity to GT12 over GT10.2. Multiple 10E8-class antibodies previously induced by vaccination with 10E8-GT nanoparticles were selected for cryoEM, and three off-target antibodies were included to help cryoEM data processing and image alignment. As expected, most 10E8-class mAbs showed higher affinities to their respective priming immunogen, likely due to the differences in the MPER binding pocket between GT10.2 and GT12. Meanwhile, the off-target mAbs W6-10 and W6-03 that recognize a conserved epitope on the scaffold showed comparable affinities to GT10.2 and GT12. Through MD simulation analysis, we observed that high affinity binders generally have more stable HCDR3 sidechain positions, while the aromatic sidechains of lower-affinity binders showed increased flexibility. Fine-tuning of the MPER binding pocket allows GT12 to better accommodate such flexibility (Fig. 1F), thus GT12 is expected to prime a broader repertoire of 10E8-like precursors than GT10.2.

Although we do not anticipate the MPER-GT-induced antibodies to bind the wild-type MPER peptide, W3-02 and W3-05 showed binding to 10E8-B1, an epitope scaffold with an almost fully native MPER sequence (except for W680N required for solubility) ^27^. The structure characterization of these 10E8-like antibodies offers important insights on the design and optimization of boosting antigens against MPER epitope and also helps in interpreting the results of future immunization studies. Previous findings suggested the double proline (PP) motif in the junction of D and J genes of HCDR3 to be important for the broad neutralizing ability of 10E8 ^27^, the potential HCDR2 interactions as well as the compatibility of LC with membrane lipid are also crucial for the correct binding angle to the MPER epitope^26,41^. However, these features are not observed in the structures of 10E8-like Fabs characterized in this study, and the priming immunogens are not designed to induce such features. To further improve the binding affinities to membrane-bound wild-type Env protein, the HCDR3 interactions need to be modified to avoid clashing with the membrane lipids and GP41, while additional interactions with paratopes outside the HCDR3 will be need to be included in boosting immunogens. Our structural studies provide a starting point to understand how potential bnAb precursors evolve to overcome undesired binding angles and avoid clashing with membrane lipid, after boosting with immunogens that are structurally closer to the full length Env.

We also mapped the off-target responses of 19 mAbs from GT10.2 12mer or GT12 12mer immunized mice using a panel of variant scaffolds with patches of mutants on their surface. The binding of two most dominant off-target epitopes were validated by cryoEM. The inclusion of multiple glycosylation sites on the scaffold was intended to mitigate off-target responses. However, previous glycan analysis of GT10.2 and GT12 nanoparticles showed 4 out of 7 designed glycosylation site on the epitope scaffolds to be poorly occupied ^27^. Here, we observed consistent presence of N-linked glycan density only on position N74/75 in the maps of monomeric GT10.2/GT12, which matches with previously reported glycosylation profiles in nanoparticles. However, we did not observe clear glycan densities at positions N3/4 and N157/158, presumably because they are located on relatively flexible N/C-termini of the monomeric scaffolds. Anecdotally, the N74/75 glycan appears to be an important feature that facilitates particle alignment during cryoEM data processing, especially for “pseudo-symmetric” complexes where two Fabs bind to the opposite sides on the scaffold. These findings underscore the need for further modifications to improve glycan occupancy, through screening alternative glycosylation positions or refining the local sequon context.

In addition to the structural analysis, we further validated the feasibility of cryoEM epitope mapping on small antigens using both homogeneous (single mAb) and heterogeneous (mAb pools) samples, suggesting it is possible to apply this approach to polyclonal Fabs. However, challenges were associated with small particle size and increased sample heterogeneity, which require optimizations during sample preparation and data processing. Here, we implemented several strategies we have tried in this work to address these problems: 1). Addition of Fabs/nanobodies or introducing elbow mutations to improve particle alignment ^42,43^; 2). Purification of complexes by size exclusion chromatography to remove unbound or weakly-bound Fabs and improve sample quality; 3). Use of gold grids to reduce movement and improve contrast during data collection; 4). Tilted-stage data collection to mitigate orientation bias; 5). Use of Topaz picker and customized CryoSPARC workflows for small particles are beneficial for particle classification.

In summary, our study provided critical structural insights for the design and optimization of HIV MPER-germline targeting vaccines. Furthermore, it presents a proof of concept for using small antigens in cryoEM-based epitope mapping. One limitation of this study is that a mixture of monoclonal antibodies might not be comparable to the actual antibody diversity in the serum. Future efforts will aim to address this problem with polyclonal serum samples.

## Methods

### Protein expression and purification

Genes encoding MPER-GT proteins and antibodies were synthesized and cloned into pHLSec or its variant pCWSec by GenScript using codons optimized for expression in human cells. Proteins were expressed and purified as described in detail previously ^27^. Briefly, plasmids were transfected into FreeStyle 293F cells (Thermo Scientific), and expression was performed in protein-free chemically defined FreeStyle medium (Thermo Scientific). His-tagged proteins were purified from clarified supernatants using immobilized metal affinity chromatography followed by size-exclusion chromatography (SEC). Antibodies were purified by protein A affinity chromatography followed by buffer exchange into Tris-buffered saline.

### Antibody isolation and characterization

The isolation and characterization of the antibodies structurally characterized in this paper was previously reported ^27^. Briefly, frozen splenocytes and lymphocytes from immunized animals were thawed subjected to B cell isolation. Streptavidin-conjugated baits were prepared by combining biotinylated monomeric baits with fluorescent streptavidin at room temperature for at least 1 h in the dark. Wild-type baits were complexed with streptavidin-AF647 (Invitrogen, S21374) and streptavidin-BV421 (BioLegend, 405225). KO baits were conjugated with TotalSeq-C hashtagged streptavidin-PE (BioLegend, 405261). Baits were conjugated with streptavidin at a 4:1 (bait/streptavidin) ratio and used at a final bait concentration of 200 nM for staining.

All samples were sorted on a BD FACSMelody using FACSChorus software. Single-color compensations were performed using C57BL/6 splenocytes with matched antibodies. For channels used for bait detection, cells were stained with biotinylated anti-CD19 (BioLegend, 115503), followed by secondary staining with the appropriate fluorescent streptavidin. Sorting gates were set to identify class switched IgG B cells that were double positive for the monomeric WT antigen and epitope specificity was determined bioinformatically by hashtag counts of the KO antigen.

Sorted samples were prepared for BCR sequencing by the 10x Genomics Single Cell Immune Profiling platform. Samples were processed according to manufacturer’s user guide for Chromium Next GEM Single Cell 5′ Reagent kits v2 (Dual Index) with Feature Barcoding. Pooled libraries were sequenced on an Illumina NextSeq 2000 using NextSeq 1000/2000 Control Software and a 100-cycle P3 reagent kit (Illumina, 20040559) with a target depth of 5,000 paired-end reads for both the V(D)J and Feature Barcode Libraries and run using read parameters indicated in the 10x Genomics user guide.

Raw sequencing data were demultiplexed, processed into assembled V(D)J contigs and counts matrix files and assigned to specific animal IDs based on TotalSeq-C antibody hashtag counts using Cell Ranger (v6.1) and scab, as previously described ^27,44^. Epitope-KO-positive cells were identified based on unique molecular identifier counts of the hashtagged PE-streptavidin probe using scab. Gene assignment, annotation and formatting into for paired heavy and light chain antibody sequences were performed using SADIE with a custom NHP or hD3-3/JH6 mouse germline reference database ^8^.

### Binding affinity determination

All *K*_d_ values for antibody–antigen interactions presented in main text figures were measured on a Carterra LSA instrument using HC30M or CMDP sensor chips (Carterra) and 1× HBS-EP+ (pH 7.4) running buffer (20× stock from Teknova, H8022) supplemented with bovine serum albumin at 1 mg ml^−1^. Carterra Navigator software instructions were followed to prepare chip surfaces for ligand capture. In a typical experiment, approximately 2,500 to 2,700 RU of capture antibody (SouthernBiotech, 2047-01) at 25 µg ml^− 1^ in 10 mM sodium acetate (pH 4.5) was amine coupled using a commercial Amine Coupling kit (GE, BR-1000-50) but using tenfold diluted NHS and EDC concentrations. Regeneration solution was 1.7% phosphoric acid injected three times for 60 s per each cycle. The solution concentration of ligands was around 1 µg ml^−1^, and contact time was 5 min. Raw sensograms were analyzed using Carterra Kinetics software (Carterra), interspot and blank double referencing, Langmuir model. For fast off-rates (>0.009 s^− 1^), we used automated batch referencing that included overlay *y* aline and higher analyte concentrations. For slow off-rates (≤0.009 s^−1^), we used manual process referencing that included serial *y* aline and lower analyte concentrations. After automated data analysis by Kinetics software, a custom R script was used to remove datasets with maximum response signals smaller than signals from negative controls.

### Epitope mapping of off-target mAbs

The antibodies were epitope mapped using a sandwich style biolayer interferometry (BLI) assay on an Octet RED384 system. All proteins were prepared in HBS-EP supplemented with 0.1% BSA. The mature 10E8 antibody was immobilized on AHC tips at a concentration of 3 ug/mL for 90 seconds, followed by a baseline in buffer of 30 seconds. The tips were then used to capture 10E8-GT10.2 or variants of GT10.2 with surface patches mutated; antigen concentration (500nM) and association time (120s) were chosen to saturate the captured 10E8 antibody with antigen. Finally, the tips were dipped into 15ug/mL off-target Fabs for 120 seconds. Data was processed on Octet Data Analysis software, version 11.1.2.24 Change in maximum nm displacement was calculated for each Fab after association with the antigen and binding was subsequently normalized by the 10E8-GT10.2 displacement. A threshold cut-off of 0.33 was used to determine loss of binding. Surface epitope specificity was determined by calculating the % binding (max nm displacement in BLI) of the elicited antibodies to patch mutants compared to WT. If there was <1/3 of the binding to the patch mutant compared to WT, the antibody was binned to that location.

### CryoEM sample preparation

The monomeric MPER-GT immunogens were incubated with the 10E8-class on-target and off-target Fabs in an equal molar ratio (1:1:1) overnight at room temperature. The complex was then purified over a Superdex 200 Increase column (GE Healthcare) and concentrated to 4.4 mg/ml (GT10.2+3 on-target Fabs+W6-03), 4.3 mg/mL (GT10.2 + 6 on-target Fabs +3 off-target Fabs), and 2.9 mg/mL (GT12+10E8+W6-10) respectively. Next, 3 μl of the complex was mixed with 0.5 μl of 35 μM lauryl maltose neopentyl glycol (Anatrace; final concentration of 5 µM) before deposition onto glow discharged 1.2/1.3 UltrAuFoil 300 grids (EMS). The grids were vitrified in liquid ethane using a Vitrobot Mark IV (Thermo Fisher Scientific) with the following settings: 4 °C, 100% humidity, 10-s wait time, 6-s blot time and blot force of 2.

### CryoEM data collection and processing

The non-tilted movie frames were collected using EPU image acquisition software (Thermo Fisher Scientific) at a nominal magnification of ×190,000 with a ThermoFisher Scientific Falcon 4 detector mounted on a ThermoFisher Scientific Glacios operating at 200 kV. Counting mode was used for data collection, with a total approximate exposure dose of 50 e^−^/Å^2^. Raw micrographs were processed with CrypoSPARC Live. The tilted dataset of SA2911+GT12+W6-10 was collected by FEI Titan Krios (Thermo Fisher Scientific) operating at 300 kV with a K2 Summit direct electron detector camera (Gatan) at a magnification of ×105,000. Data was automated collected using the Leginon software package ^45^ at 30° tilt angle with a total exposure dose of 52.6 e^−^ /Å^2^. Image pre-processing was performed with the Appion software package ^46^, and tilted frames were aligned, dose-weighted using UCSF MotionCor2 software ^47^, and GCTF was estimated ^48^. All preprocessed micrographs were used for blob picking and 2D classifications in CryoSPARC v4.0, initial high quality 2D classes were further used for template picker or to train Topaz picker ^49^. Additional information can be found in Table S1 and Figure S2.

### CryoEM model building

For model building, an initial model of 10E8 Fab and MPER scaffold was generated using PDB 8SX3 and docked into the cryo-EM map using UCSF ChimeraX^50^. For the other on-target and off-target antibodies, the initial Fv models were generated by SAbPred ABodyBuilder2 ^51^ and docked into the cryoEM maps. Coot 0.9.8 ^52^, Phenix^53,54^ and Rosetta ^55,56^ were used for model building and refinement. The final models and maps have been deposited in the PDB and Electron Microscopy Data Bank under accession codes listed in Table S1.

### Molecular Dynamics (MD) simulations

As starting structures for our MD simulations, we used the available cryo-EM structures of GT10.2 and in complex with 10E8 and W3-05 and GT12 in complex with 10E8 and SA2911 (this study). The additional variants GT10.2 in complex with SA2911 and GT12 in complex with W3-05 were modelled using the respective cryo-EM structures as templates. The starting structures for our simulations were prepared in MOE (*Molecular Operating Environment*, 2022.02 Chemical Computing Group ULC, 910-1010 Sherbrooke St. W., Montreal, QC H3A 2R7, 2022) using the Protonate3D tool (MOE ^57^)

To neutralize the charges, the uniform background charge was applied, which is required to compute long-range electrostatic interactions ^58^. Using the tleap tool of the AmberTools22 ^59^ package the structures were soaked in cubic water boxes of TIP3P water molecules with a minimum wall distance of 12 Å to the protein ^60–62^. For all simulations, parameters of the AMBER force field 14SB were used ^63^.

We then performed two repetitions of 1 µs of classical molecular dynamics simulations for each variant, resulting in an aggregated simulation time of 12 µs of simulation time. Molecular dynamics simulations were performed in an NpT ensemble using pmemd.cuda ^64^. Bonds involving hydrogen atoms were restrained by applying the SHAKE algorithm ^65^ allowing a time step of 2 fs.

The Langevin thermostat was used to maintain the temperature during simulations at 300 K ^66,67^ with a collision frequency of 2 ps^−1^ and a Monte Carlo barostat ^68^ with one volume change attempt per 100 steps.

The interaction energies were calculated with cpptraj using the interaction energy (LIE) tool ^69^. The electrostatic interaction energies were calculated for all frames of each simulation and provided the simulation-averages of these interactions. Additionally, we performed a hierarchical agglomerative clustering implemented in cpptraj, clustering on the Cα-positions all CDR loops using a distance cut-off criterion of 3 Å. PyMOL was used for visualizing protein structures (The PyMOL Molecular Graphics System, Version 2.5.2 Schrödinger, LLC).

## RESOURCE AVAILABILITY

### Lead contact

ABW

### Materials availability

Plasmids or proteins related to the immunogens, sort reagents, or antibodies employed in this study are available from WRS or TS under a material transfer agreement with The Scripps Research Institute.

### Data and code availability

Cryo-EM atomic models have been deposited in the Protein Data Bank with accession codes 9PV3, 9PV4, 9PV5 (Table S1). All cryo-EM maps are available in the Electron Microscopy Data Bank (EMDB) under accession codes EMD-71881, EMD-71882, EMD-71883, EMD-71884, EMD-71885 (Table S1).

## ACKNOWLEDGMENTS

We thank Hannah Turner, Anant Gharpure, and Will Lessin for their support on electron microscopy, Charles Bowman and J.C. Ducom for compute support. This work was supported by National Institute of Allergy and Infectious Diseases (NIAID) UM1 Al100663 (Scripps Center for HIV/AIDS Vaccine Immunology and Immunogen Discovery, CHAVI-ID), UM1 AI144462 (Scripps Consortium for HIV/AIDS Vaccine Development, CHAVD) (to TS, WRS, and ABW), R01 AI113867 (to TS and WRS), the Bill and Melinda Gates Foundation Collaboration for AIDS Vaccine Discovery (NAC INV-007522, INV-008813, and INV-034657 to WRS and ABW), IAVI support to WRS, and the IAVI Neutralizing Antibody Center (NAC) to WRS and ABW. OMS is supported in part by the National Institute of Allergy and Infectious Diseases (NIAID) grant number 5F31AI179426-02.

## AUTHOR CONTRIBUTIONS

Conceptualization, JH, OS, KR, GO, TS, WRS, ABW

methodology, JH, OS, KR, MFQ, JL, RT, EG, NP, GO, TS, WRS, ABW

Investigation, JH, OS, KR, MFQ, JL, RT, EG, NP, GO, TS

writing—original draft, JH, OS, KR

writing—review & editing, JH, OS, KR, MFQ, JL, RT, EG, NP, GO, TS, WRS, ABW

funding acquisition, TS, WRS, ABW

resources, JH, OS, KR, MFQ, JL, RT, EG, NP, GO, TS, WRS, ABW

supervision, TS, WRS, ABW

