## Supplemental Information for "CryoEM Structures of Antibodies Elicited by Germline-Targeting HIV MPER Epitope-Scaffolds"

### Table S1. CryoEM data collection, processing and model building statistics

| Model/Map | GT10.2+W3-05+W6-03 | GT10.2+W3-02+W6-03 | GT10.2+W10-09+W6-10 | GT12+10E8+W6-10 | GT12+SA2911 + W6-10 |
|---|---|---|---|---|---|
| **EMDB** | EMD-71883 | EMD-71885 | EMD-71884 | EMD-71882 | EMD-71881 |
| **PDB** | 9PV5 | - | - | 9PV4 | 9PV3 |
| **Data collection & processing** | | | | | |
| **Microscope/ Detector** | Glacios/Falcon 4 | Glacios/Falcon 4 | Glacios/Falcon 4 | Glacios/Falcon 4 | Krios/Gatan K3 |
| **Voltage (kV)** | 200 | 200 | 200 | 200 | 300 |
| **Magnification** | 190kx | 190kx | 190kx | 190kx | 105kx |
| **Recording mode** | Counting | Counting | Counting | Counting | Counting |
| **Pixel size (Å)** | 0.725 | 0.725 | 0.725 | 0.725 | 0.833 |
| **Total dose ($e^-/Å^2$)** | 50.4 | 50.4 | 46.5 | 43.2 | 52.6 |
| **Defocus range (µm)** | -0.7 to -1.5 | | -0.8 to -1.8 | -0.8 to -1.8 | -1.0 to -2.0 |
| **No. of movie micrographs** | 6090 | | 6918 | 3585 | 4104 |
| **No. of molecular projection images in map** | 41,951 | 63,258 | 12,994 | 72,061 | 171,994 |
| **Symmetry** | C1 | C1 | C1 | C1 | C1 |
| **Map pixel size (Å)** | 0.725 | 0.725 | 0.725 | 0.725 | 0.833 |
| **Map resolution (FSC 0.143; Å)** | 3.93 | 4.21 | 6.31 | 4.01 | 3.95 |
| **Map sharpening B-factor (Å2)** | -101.8 | -162.3 | -490.3 | -128.2 | -170.3 |
| **Structure building and validation** | | | | | |
| **Model composition** | | | | | |
| **Non-hydrogen atoms** | 4963 | - | - | 4831 | 4862 |
| **Protein residues** | 630 | - | - | 616 | 618 |
| **Ligands** | NAG:1 | - | - | NAG:1 | NAG:1 |
| **RMSD bond length (Å)/angles (°)** | 0.008/1.459 | - | - | 0.018/1.749 | 0.008/1.411 |
| **MolProbity score** | 1.63 | - | - | 1.24 | 1.72 |
| **EMRinger score** | 1.05 | - | - | 2.89 | 2.1 |
| **Clash score** | 4.92 | - | - | 3.05 | 5.64 |
| **Ramachandran outliers/allowed/favored (%)** | 0/5.48/94.52 | - | - | 0/2.81/97.19 | 0/6.25/93.75 |
| **Rotamer outliers (%)** | 0 | - | - | 0 | 0.19 |
| **Cβ outliers (%)** | 0 | - | - | 0 | 0 |

**Table S2. Off-target epitope mapping with patch mutations of GT10.2**

| % of GT10v55 binding | | | | | | | |
|---|---|---|---|---|---|---|---|
| | **Patch 1** | **Patch 2** | **Patch 3** | **Patch 6** | **Patch 7** | **Patch 8** | **Patch 9** |
| **SA684_Exp1_Scaf_MPER-GT10v55_W6-01** | 1.13 | 0.26 | 1.10 | 0.96 | 1.09 | 0.87 | 1.09 |
| **SA684_Exp1_Scaf_MPER-GT10v55_W6-02** | 0.93 | 0.94 | 0.73 | 0.77 | 0.94 | 0.60 | 0.62 |
| **SA684_Exp1_Scaf_MPER-GT10v55_W6-03** | 0.90 | 0.88 | 0.89 | 0.90 | 0.92 | 0.33 | 0.28 |
| **SA684_Exp1_Scaf_MPER-GT10v55_W6-05** | 1.29 | 0.52 | 1.18 | 0.76 | 1.32 | 0.98 | 1.16 |
| **SA684_Exp1_Scaf_MPER-GT10v55_W6-06** | 0.88 | 0.96 | 0.78 | 1.00 | 0.80 | 0.24 | 0.26 |
| **SA684_Exp1_Scaf_MPER-GT10v55_W6-07** | 0.83 | 0.95 | 0.79 | 1.02 | 0.86 | 0.58 | 0.87 |
| **SA684_Exp1_Scaf_MPER-GT10v55_W6-08** | 0.84 | 0.88 | 0.94 | 0.93 | 0.96 | 0.94 | 0.90 |
| **SA684_Exp1_Scaf_MPER-GT10v55_W6-09** | 1.22 | 0.74 | 1.29 | 0.94 | 1.12 | 0.83 | 1.15 |
| **SA684_Exp1_Scaf_MPER-GT10v55_W6-10** | 0.96 | 0.88 | 0.99 | 0.94 | 1.02 | 0.75 | 0.73 |
| **SA684_Exp1_Scaf_MPER-GT12v56_W6-01** | 0.68 | 0.78 | 0.76 | 0.80 | 0.79 | 0.66 | 0.48 |
| **SA684_Exp1_Scaf_MPER-GT12v56_W6-02** | 0.93 | 0.72 | 0.98 | 0.81 | 1.03 | 0.95 | 0.92 |
| **SA684_Exp1_Scaf_MPER-GT12v56_W6-03** | 0.85 | 0.97 | 0.80 | 0.99 | 0.73 | 0.81 | 0.77 |
| **SA684_Exp1_Scaf_MPER-GT12v56_W6-04** | 0.90 | 0.39 | 0.84 | 0.96 | 0.93 | 0.99 | 0.90 |
| **SA684_Exp1_Scaf_MPER-GT12v56_W6-05** | 0.86 | 0.94 | 0.92 | 0.96 | 0.95 | 0.69 | 0.95 |
| **SA684_Exp1_Scaf_MPER-GT12v56_W6-06** | 1.29 | 0.71 | 1.38 | 0.89 | 1.19 | 0.77 | 1.24 |
| **SA684_Exp1_Scaf_MPER-GT12v56_W6-07** | 0.97 | 0.88 | 0.94 | 0.95 | 0.99 | 0.19 | 0.25 |
| **SA684_Exp1_Scaf_MPER-GT12v56_W6-08** | 0.96 | 0.80 | 0.43 | 0.87 | 1.05 | 0.78 | 0.91 |
| **SA684_Exp1_Scaf_MPER-GT12v56_W6-09** | 0.96 | 0.16 | 0.92 | 0.90 | 0.98 | 0.64 | 0.36 |
| **SA684_Exp1_Scaf_MPER-GT12v56_W6-10** | 0.90 | 0.95 | 0.81 | 0.98 | 0.88 | 0.27 | 0.46 |
| **10E8UCA** | 0.87 | 0.84 | 0.53 | 0.99 | 0.90 | 1.00 | 0.87 |

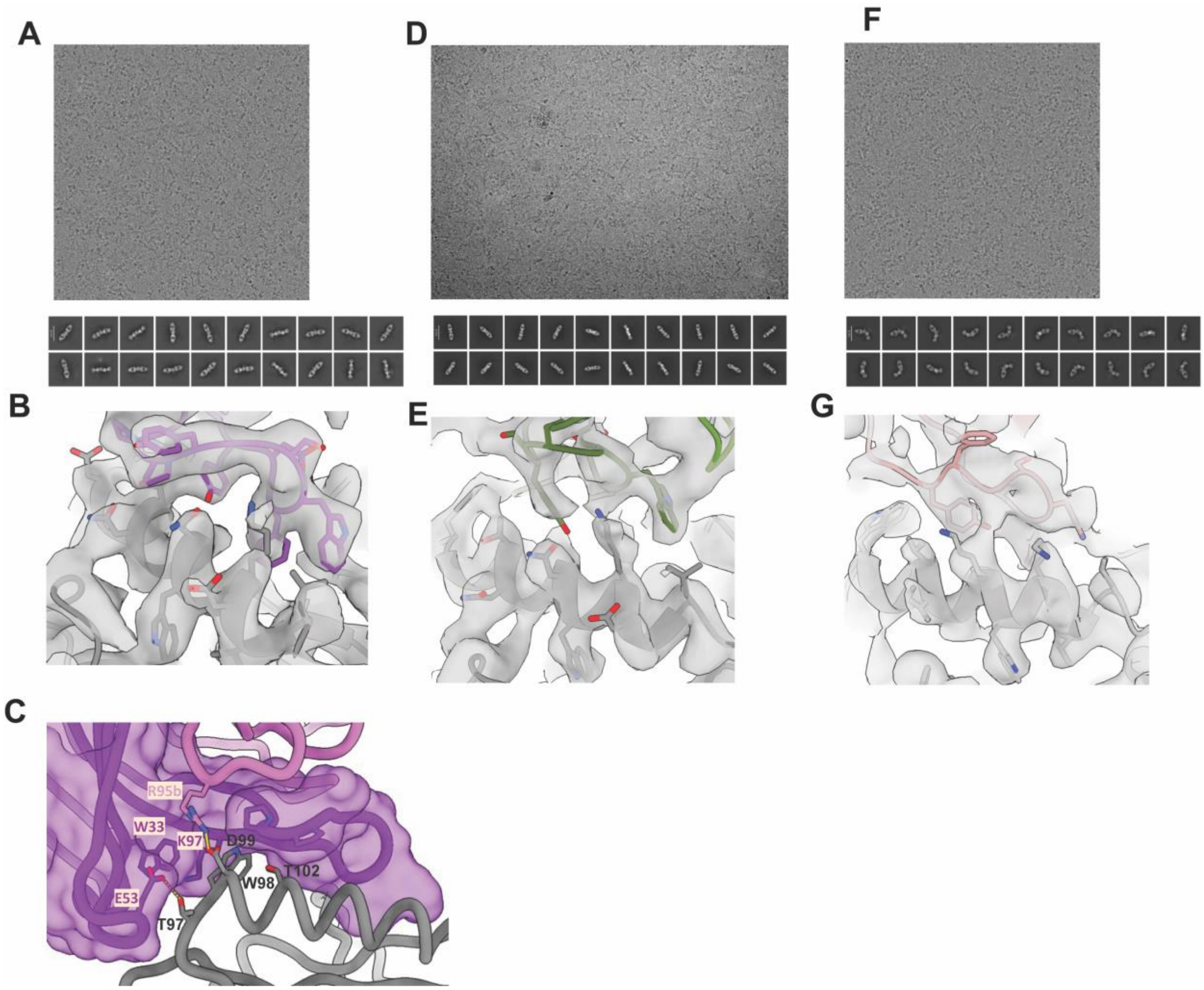


**Figure S1.** Representative micrograph, selected 2D classes, and density map near MPER helix are shown for 10E8+GT12+W6-10 complex (A & B), SA2911+GT12+W6-10 complex (D & E), and W3-05+GT10.2+W6-03 complex (F & G). (C). GT12 interactions with 10E8 (non-HCDR3 mediated) with 10E8 HC shown with surface.

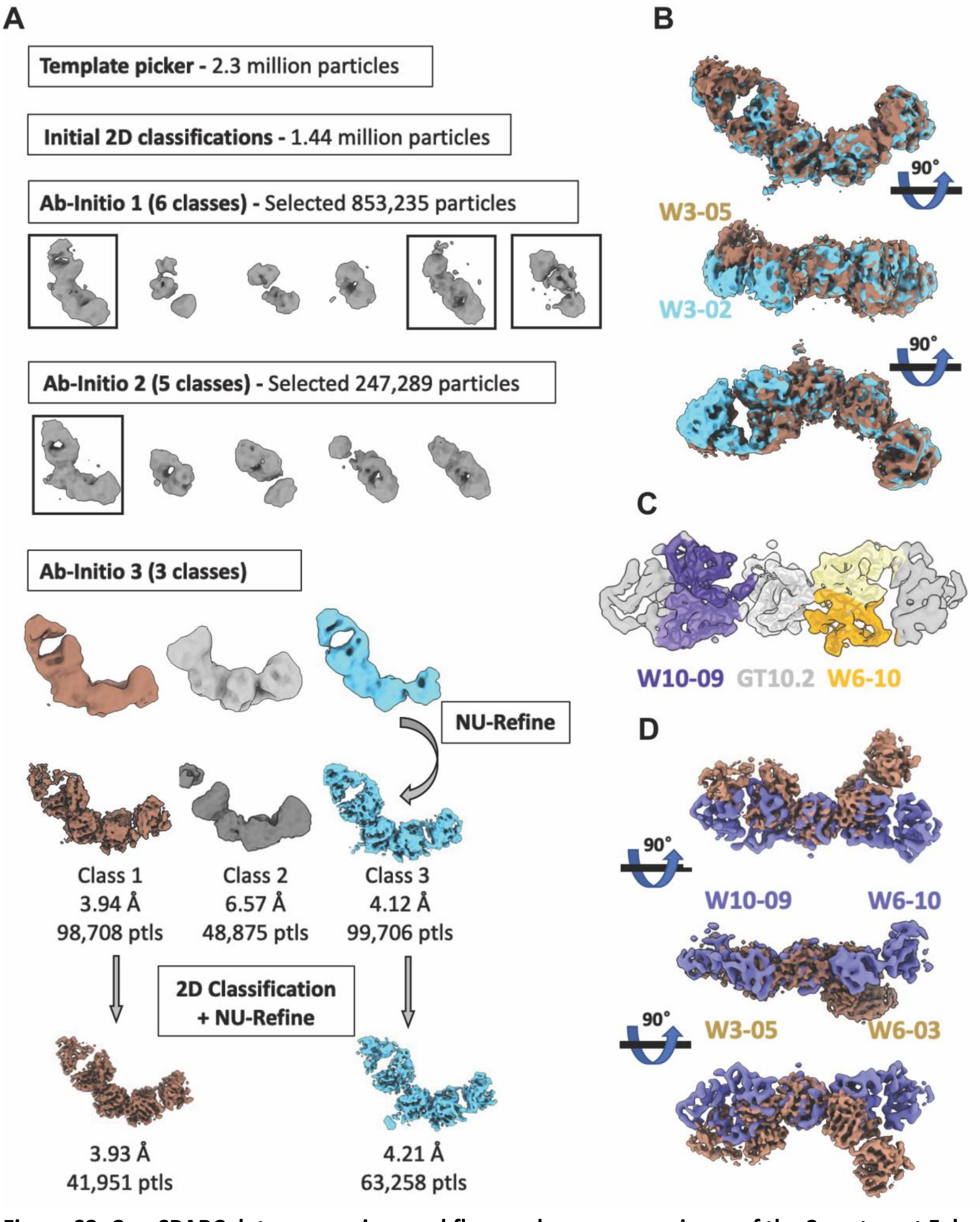


**Figure S2. CryoSPARC data processing workflow and map comparisons of the 3 on-target Fab pool dataset.** (A). Particles were cleaned up and classified through 2D classification and multiple rounds of Ab-initio jobs and final maps were obtained by non-uniform refinement jobs. (B). Aligned maps from 2 different classes showing the shift of binding angle of the on-target

Fabs. (C). CryoEM map of W10-09 (purple)/W6-10 (yellow) Fabs complexing with GT10.2. (D). Aligned maps of W3-05 + GT10.2 + W6-03 vs. W10-09 + GT10.2 + W6-10 showing different binding angles of on/off-target Fabs.

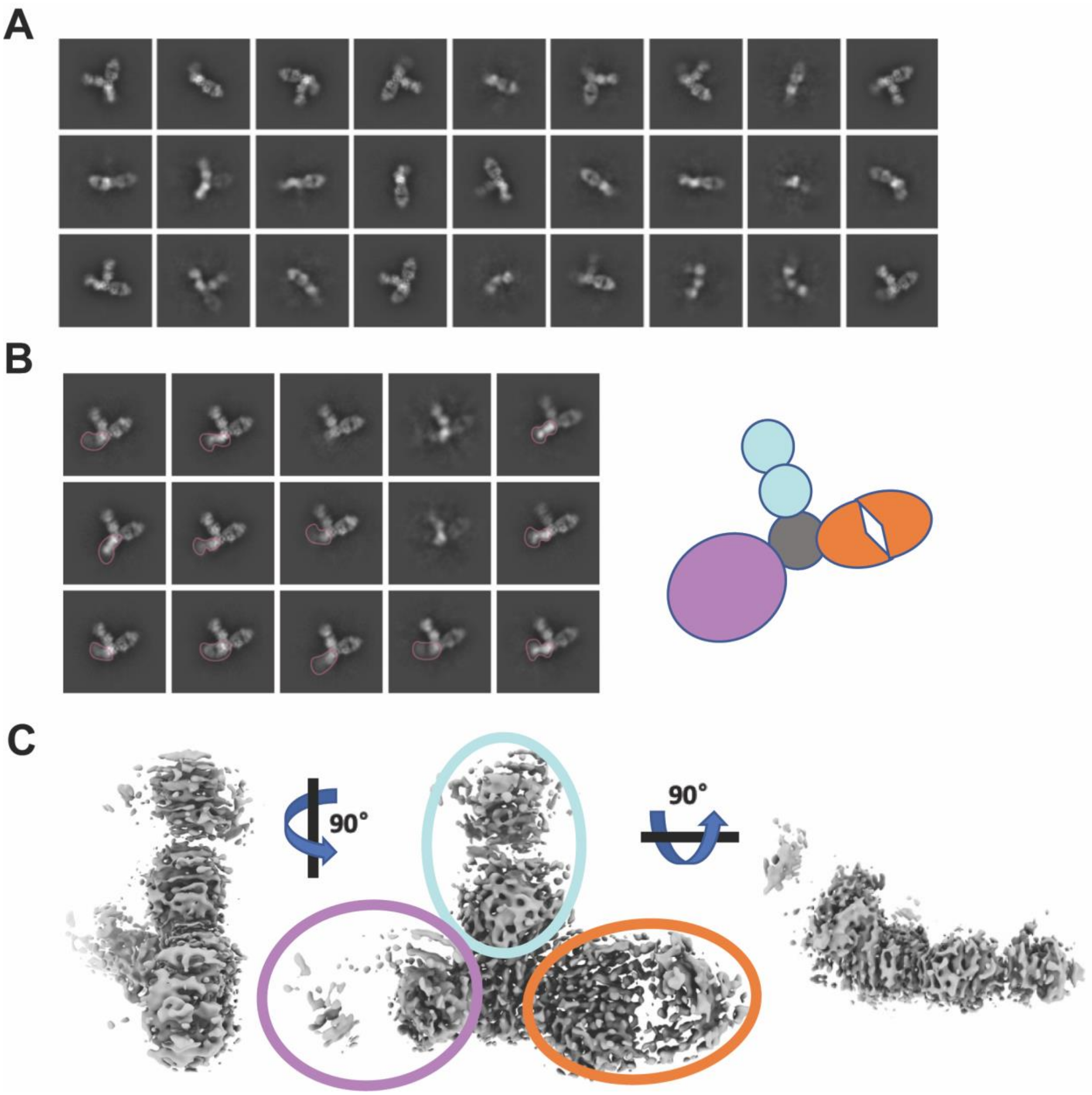


**Figure S3. Selected 2D classes and map from 6 on-target Fab pool dataset.** Selected 2D classes (A) and representative superclass from a CryoSPARC rebalance 2D classes job (B) showing a variety of binding poses on on-target Fabs (circled in purple) from the pool. (C). A 3D reconstruction map showing Fab densities on 3 different binding sites of HGT10.2 immunogen.

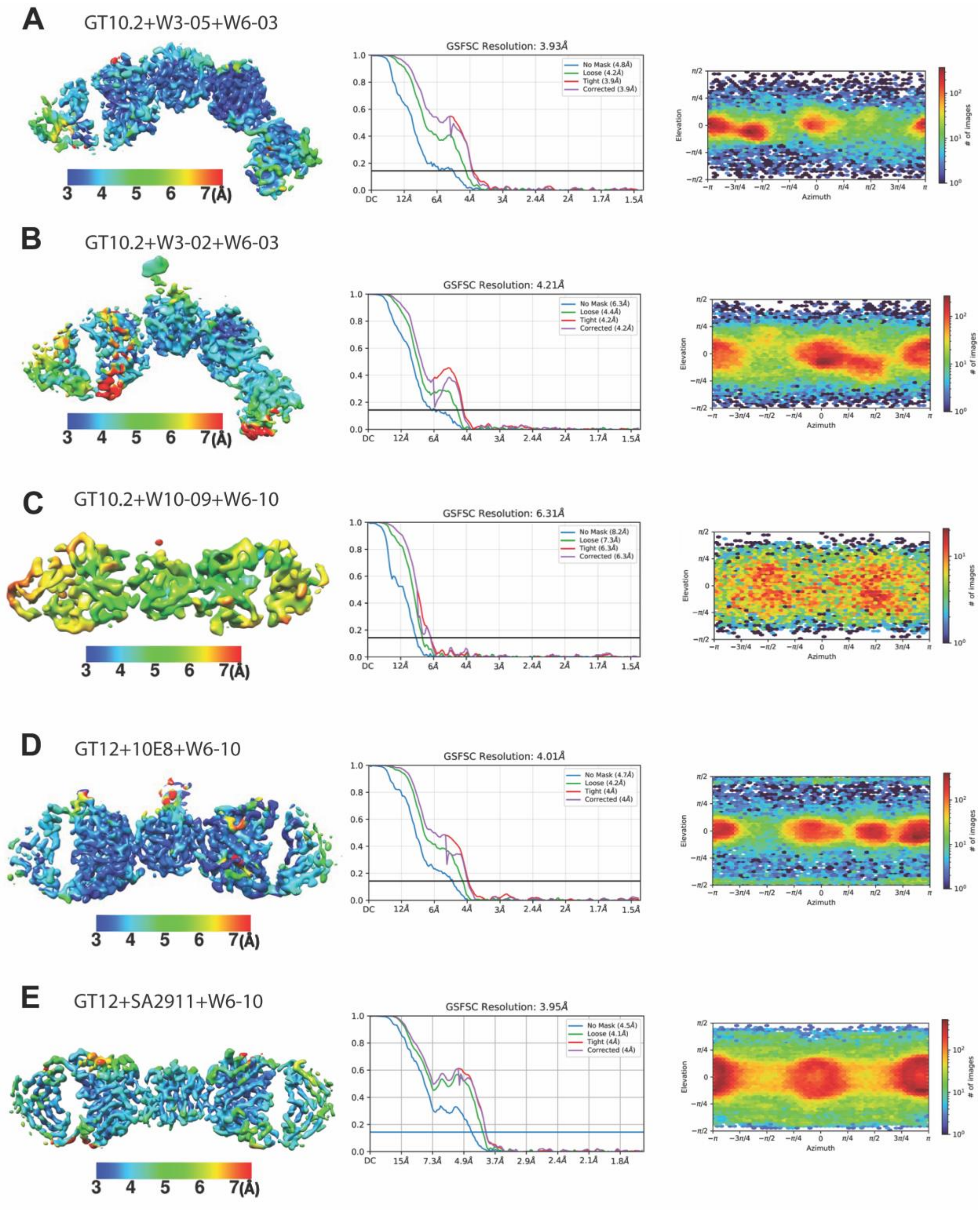


**Figure S4. The local resolution maps, Fourier shell correlation curves and angular sampling for cryoEM maps in this manuscript.**

**Video 1. MD simulation of 10E8+GT10.2**

**Video 2. MD simulation of 10E8+GT12**

**Video 3. MD simulation of W3-05+GT10.2**

**Video 4. MD simulation of SA2911+GT12**
